# A critical affinity window for IgSF proteins DIP-α and Dpr10 is required for proper motor neuron arborization

**DOI:** 10.1101/2024.09.11.612484

**Authors:** Davys H. Lopez, Kevin D. Rostam, Sumaira Zamurrad, Shuwa Xu, Richard S. Mann

## Abstract

For flies to walk properly, motor neurons (MNs) from the ventral nerve cord (VNC) need to reach the correct muscle, and arborize appropriately during development. The canonical view of how this is achieved is that cell surface proteins are expressed pre- and post-synaptically that bind to each other like molecular “lock-and-keys” that guide neurons to their targets. The binding affinities of these molecules can vary by more than 100-fold. In the fly leg neuromuscular system, three MNs express *DIP-α* and their target muscles express its cognate partner, *dpr10*, both of which encode members of the Immunoglobulin superfamily (IgSF). Although, both of these molecules are necessary for the maintenance of MN-muscle contacts, the role that specific affinities play in this process has not been examined. Here we use knock-in mutations into *DIP-α* and *dpr10* that either decrease or increase the affinity between these two proteins. Compared to control animals, decreasing the affinity results in phenotypes similar to *DIP-α* or *dpr10 n*ull animals, where MN axons fail to maintain contacts with their muscle targets and retract their filopodia, resulting in stunted and/or branchless axons. We also find that the three *DIP-α*-expressing motor neurons behave differently to the loss of affinity. Surprisingly, if the affinity increases past a certain threshold, a similar branchless phenotype is observed in adult legs. Live imaging during pupal development shows that MN filopodia are unable to productively engage their muscle targets and behavioral assays suggest that these defects lead to locomotor deficits. These data suggest that CAM affinities are tuned to a specific range to achieve proper neuronal morphology.

## Introduction

Animal movement requires developing motor neurons (MNs) to arborize and synapse onto the correct muscles. For axons to establish the precise set of synapses required for neural circuit formation, it is helpful to conceptualize this process into a series of broad steps: axon guidance or pathfinding, in which axons navigate close to their cellular targets^1–4^, synaptic specificity, where short-range interactions establish synaptic choice^2,5,6^, branching and arborization^7,8^, and synaptic plasticity, which is an activity-dependent process that can modify preexisting synapses^9,10^. In all these steps, cell adhesion molecules (CAMs) have been implicated as playing critical roles^2,5^. For example, CAMs that bind hetero- and homophilically such as the *beat-Ia* and *sidestep* have been implemented during axon pathfinding^3,4,11,12^, molecules such as the L1 protein subfamily are needed for synaptic specificity^5,13–15^, Dscam1 and Protocadherins are needed for self-avoidance^16–18^ and β1 integrins and contactins are used for synaptic plasticity^5,9,10,17,19–21^

In Drosophila, the DIPs (Dpr-interacting proteins) and Dprs (Defective proboscis response) are a subset of CAMs that are also members of the Ig domain Superfamily (IgSF) proteins. The “Dpr-ome” interaction network was initially discovered using an *in vitro* interactome assay that detected interactions between extracellular protein domains^22,23^. This interactome and its subsequent analysis revealed that there are 9 Dprs, each with two Ig domains, that bind selectively to DIPs, which have 21 members, each with three Ig domains^22,24,25^

DIPs and dprs are expressed by a subset of neurons, suggesting a role in synaptic specificity. For instance, photoreceptors R7 and R8 and lamina neurons L1-5 in the fly visual system express a subset of Dprs, with their axonal targets in the medulla expressing matching DIPs ^24^. In the larval nervous system Dprs are more broadly expressed than DIPs in both MNs and SNs, and each neuron expresses a unique combination of Dpr and DIP genes^26^. This is also the case in the adult leg neuromuscular system where *dprs* are more widely expressed than the *DIPs*, in the majority of adult leg MNs and leg sensory neurons^27^. Furthermore, in the mushroom body, 16 of 21 *dpr* members were shown to be dynamically expressed in developing γ-KCs. Together, this indicates that DIPs and Dprs expressed by different neurons are likely to be involved in many stages of neural development^28^.

The DIPs and Dprs have also been shown to play an important role in establishing neuronal morphologies and constructing neural circuits. For example, *dpr11* and *DIP-*γ mutant animals have defects in presynaptic terminal development in the larval neuromuscular system and in the visual system both are required for the photoreceptor neuron, yR7, to target the correct M6 layer^25^. Furthermore, *DIP-*γ mutant flies also have a reduction in Dm8 neurons compared to controls^25^. *DIP-*ε has also been shown to have a critical role in establishing wildtype *fruP1* neuronal patterns, in both a cell-autonomous and non-cell-autonomous manner^29^. And, Dpr12/DIP δ transneuronal interactions are required for γ-Kenyon cells to find their proper targets in the mushroom body^28^. In the optic lobe, DIP-α heterophilic interactions with Dpr6 and 10 regulate arborization within layers, synapse number, layer specificity, and cell survival of amacrine-like Dm4 and Dm12 neurons^30^.

In the larval motor system, *DIP-α* is expressed by only two motor neurons (MNs) in each hemisegment, ISNb/d-1s and ISN-1s. In *DIP-α* mutants, one branch of the ISN-1s MN fails to innervate muscle 4 (m4)^31^. In addition, in *DIP-α* mutants, the axons that fail to innervate m4 now innervate muscle m1^31^. Interestingly, during development ISN- 1s still projects to m4 but fails to be stabilized. DIP-α cognate partners, Dpr6 and Dpr10, are expressed in 1s and in muscles, and knockdown of *dpr10* in the muscle, or in *dpr10* null animals, there is 100% loss of 1s boutons on m4 similar to the *DIP-α* null. *dpr6* hemizygous mutants have no obvious defect in ISN-1s^31^.

*DIP-α* also has a role in the adult leg motor system. Using live imaging, it has been shown that the leg neuromuscular system develops through sequential rounds of stereotyped MN branching and defasciculation followed by muscle targeting, arborization, and synapse formation^27^. Using gene trap reporters into *DIP-* ^27,32^, it was shown that it is expressed by three MNs that target the long tendon <u>m</u>uscle in the <u>fe</u>mur, αFe-ltm; the long tendon <u>m</u>uscle in the tibia, αTi-ltm; and the <u>ta</u>rsal depressor <u>m</u>uscle in the tibia, αTi-tadm (Figure 1C). In *DIP-α* null mutants, the terminal axon branches of αFe-ltm are absent in almost all samples, while those of the αTi-ltm are missing in approximately 70% and those of the αTi-tadm are rarely missing. *dpr10* is expressed broadly by all muscles and in *dpr10* null animals the three DIP-α-expressing MNs display the same phenotype as *DIP-α* null animals^27^. To differentiate between whether axons fail to send out filopodia or whether they send out filopodia but are not able to arborize, live imaging was used to visualize developing *DIP-α* null axons^27^. These videos showed that, compared to controls with an intact *DIP-α* locus, axons send out filopodia to the correct muscle target but fail to arborize and eventually retract, resulting in adults with no axon branches^27^.

**Figure 1.**
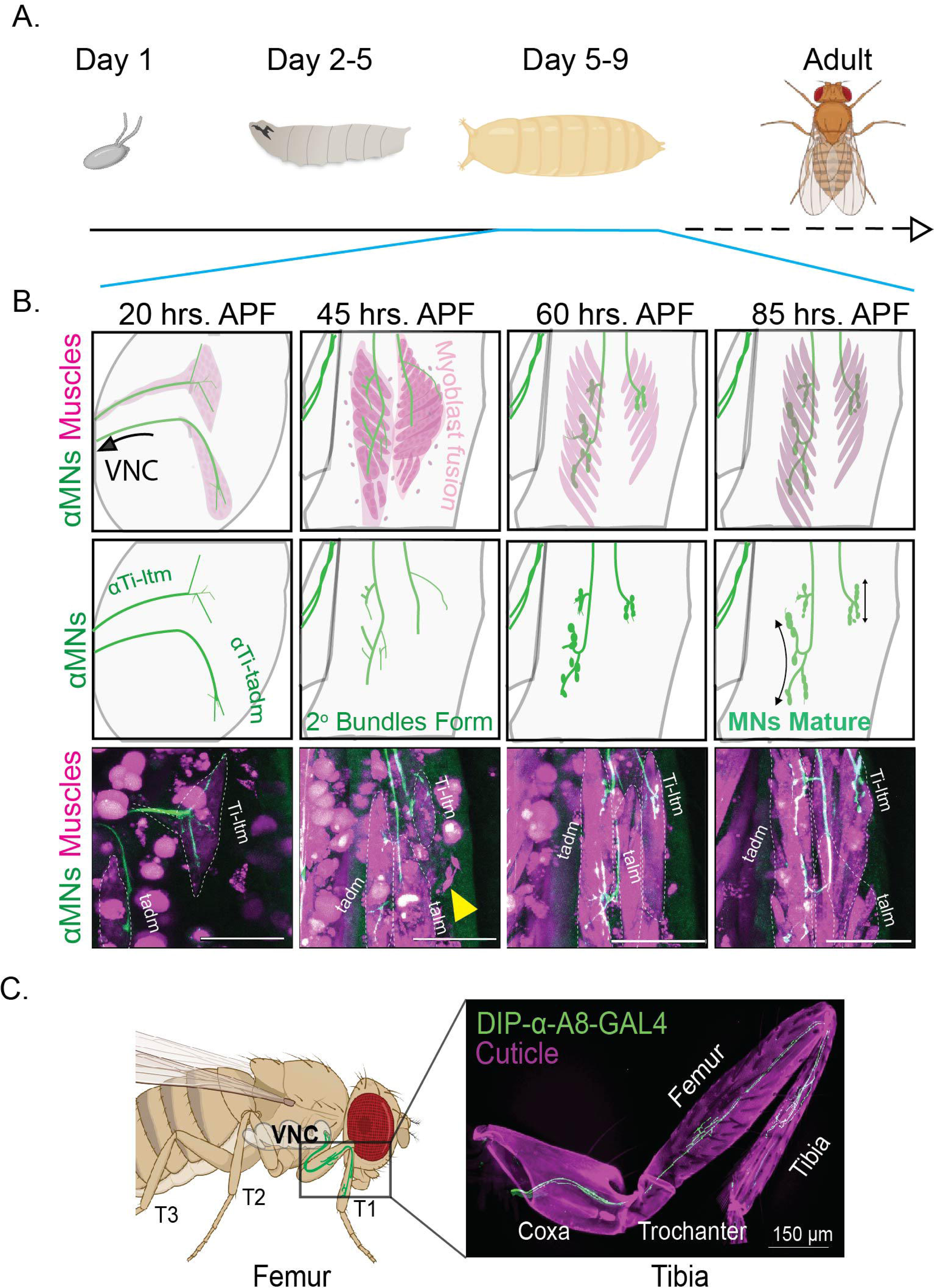
Overview of fly leg MN arborization. **A**. Life cycle of *Drosophila*. Imaging was done during most of pupal development, from 20 hours (hrs) APF and onwards. **B.** Drawings and individual frames from videos of *DIP-α-T2A-Gal4>>20X6XGFP, Mef2-QF>>10XQUAS-6XmCherry* flies. First row shows αMNs in green and muscles in magenta. Sequence shows αMNs developing in the context of myoblast fusion. Second row shows developing αMNs. At early time points, αMNs are more mobile but from 60 hrs APF onwards axon termini begin to stabilize yet continue to mature until eclosion. Third row shows stills from the video (see Video 1). **C.** Adult fly (left) and leg (right). αMNs, visualized here with *A8-Gal4, 20X6XUAS-GFP*, are born in the VNC and extend to arborize on muscles in the tibia and femur. Ti-ltm= Tibia long tendon muscle, tadm-tarsal depressor muscle, talm-tarsal levator muscle

The binding affinities of all interacting DIP/Dpr pairs have been measured using biophysical methods. K_D_s for homophilic and heterophilic binding for the DIP/Dpr pairs vary by more than 100-fold, ranging from <2 to >200 μM^24,33^. Interestingly, DIP-α and Dpr10, the binding pair examined here, have one of the highest affinity interactions in the DIP-Dpr interactome, with a K_D_ of 1.4 μM^24,33^. Motivated by these observations, a series of mutations were designed based on X-ray crystal structures that include the DIP-α–Dpr10 binding interface that either decrease or increase binding affinity, and were knocked into the *DIP-α* and *dpr10* loci^33^. In the optic lobe, it was shown that reducing the affinity of either *DIP-α* and *dpr10* causes loss-of-function phenotypes such as synaptic targeting and cell survival defects, and that the severity of these phenotypes scales with the magnitude of affinity reduction^33^. Interestingly, although increasing affinity did not alter targeting, it did affect cell survival in this system. These results indicate that affinity are tuned to achieve proper cell survival and synaptic connections^33^. In contrast to the visual system, no affect on leg MN survival has been observed in *DIP-α* and *dpr10* mutants. This may be because in the visual system, neurons are produced in excess and those that do not form functional synapses are eliminated via apoptosis^34^. In contrast, the wiring of the leg motor neuron to muscle innervation is hardwired and every individual leg motor has a unique target and morphology^35,36^.

Here, we take advantage of the precision of the leg motor neuromuscular system to examine the role of affinity between DIP-α and Dpr10. We find that decreasing or increasing affinity does not affect cell survival nor does it cause mistargeting, but rather has a role in stabilizing MN axon arborization. Using a quantitative neuron tracing analysis^37^ we find that the three DIP-α-expressing MNs (αMNs) have different affinity requirements. Furthermore, we find that increasing affinity does not result in more branches but causes aberrant neuronal morphology. Furthermore, we show that changing the affinity affects proper locomotion. Together, these results reveal that affinities between interacting CAMs have been tuned to achieve proper neuronal development.

## Results

### Leg MNs arborize through all of pupal development

We used time lapse imaging to obtain a better description for how αMNs coordinate their branching as they contact nascent groups of muscle precursors. Previously, we live imaged αMNs until about 45 hours After Pupal Formation (hrs APF)^27^. However, to extend this analysis to axon morphology changes that occur between mid-pupa and adult we improved our live imaging protocol, allowing us to visualize development later in pupal stages. We used *DIP-α-T2A-Gal4>20XUAS-6XGFP* to visualize the three αMNs together with *Mef2-QF>QUAS-10XmCher*ry to visualize muscles as they develop from ∼20-108 hrs APF, which encompasses almost all of pupal development (Figure 1A and Videos 1 and S1). Because we can only visualize structures close to the coverslip, we are limited to visualizing MNs and muscles in the tibia, which is close to the surface of the pupa.

As early as 20 hrs APF αTI-MNs have already targeted nascent clusters of myoblasts within the tibia, as previously described^27^. Interestingly, by visualizing the developing myoblasts with myr::tdTomato, we find that the myoblasts are organized into interconnected clusters that will become future muscle groups i.e. the tarsal depressor muscles (tadm) and the Tibia long tendon muscle (Ti-ltm) (Video 2 and Figure S4). These clusters then split and become discrete muscle groups. Between 25-55 hrs APF, the myoblasts will fuse and become a syncytial multifiber muscle (Figure 1B). By 20-60 hrs APF, the videos show single mobile myoblasts that end up joining and fusing with a particular muscle fiber (Figure 1B). At ∼50 hrs APF αTi-MNs have reached and stabilized a nascent axon terminus with a limited number of branches on their targeted muscle. However, from ∼50 hrs APF and onward, this initial nascent axon terminus will continue to give rise to new branches and elongate its existing branches until a few hours before the animal ecloses (Figure 1B). During this later phase, the movement of the αMN filopodia will slow and take on a bead-on-string morphology, indicating a transition from a growth cone to synapse formation (Video 1). Based on these observations, there are at least two different phases of axon morphogenesis, an initial phase where the αTi-MNs choose and stabilize muscle fibers as they are developing, and a second phase when αTi-MNs limit their mobility but continue to arborize while forming synapses. This indicates that MN arborization occurs for much longer than previously noted^27^.

### Reducing DIP-**α** affinity to Dpr10 impedes arborization

Reducing the affinity between DIP-α and Dpr10 causes cell death of Dm4 and Dm12 neurons in the optic lobe and mistargeting of Dm12 from medulla layers M3 to M8^33^. Using the same *DIP-α* and *dpr10* alleles used in that study we examined the final extent of leg MN arborization. These alleles alter the affinity between DIP-α and Dpr10 in graded fashion over a wide range, do not affect binding specificity, and do not alter homophilic binding between DIP-α and itself. We used the following *DIP-α* mutants: K81Q (K_D_ = 31.8 μM, *DIP-α*^−*20F*^; superscripts indicate the direction and fold change (F) in affinity relative to wild type), K81Q G74S (K_D_ = 68.0 μM, *DIP-α*^−*50F*^) (Figure 2A) and *dpr10* mutants: V144K (K_D_ = 11.3 μM, *dpr10*^−*8F*^), Q138D (K_D_ = 27.6 μM, *dpr10*^−*20F*^), and G99D, Q142E, V144K (K_D_ = 50.0 μM, *dpr10*^−*40F*^) (Figure 3A). Although *dpr10*, but not *dpr6,* is necessary for the terminal branching of the αMNs^27^, all *dpr10* mutant chromosomes are also mutant for *dpr6*. The αMNs were visualized using a previously characterized MiMIC line converted to T2A-Gal4, and repeated using a transcriptional enhancer isolated from *DIP-α* called A8 fused to Gal4 (*A8-Gal4*)^27^ (Figure S1). To acquire quantitative information regarding branch number and branch length, we traced and quantified both parameters using the semi-automatic tracing software on FIJI^37^, and compared control animals with wildtype alleles to animals carrying either one copy of the mutant *DIP-α* allele and one copy of the *DIP-α-T2A-Gal4* (which is a strong hypomorph^27^) or two copies of the *DIP-α* allele with *A8-Gal4* (Figure S1)^37^. We also examined animals with null alleles as a negative control.

**Figure 2.**
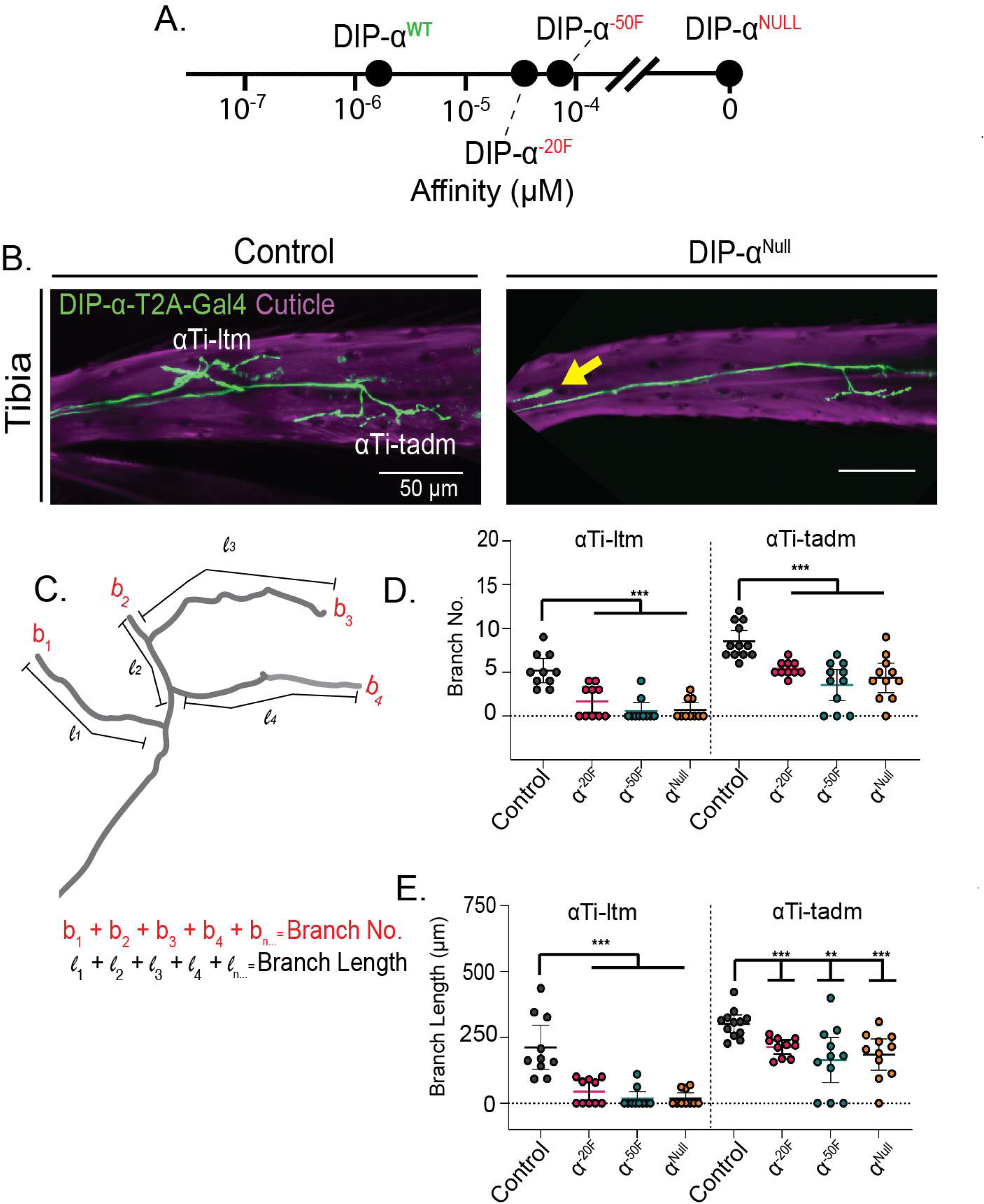
Analysis of *DIP-α* alleles with reduced affinity. **A.** Affinity range of *DIP-α* mutants in log scale. **B.** Representative images of *yw/DIP-α-T2A-Gal4>>20XUAS-6XGFP* (control) and *DIP-α^null^/DIP-α-T2A-Gal4>>20XUAS-6XGFP* tibia leg segments. **C.** Schematic of how MN morphologies are quantified. **D**. Quantification of terminal branch number of *DIP-α* mutant alleles over *DIP-α-T2A-Gal4>>20XUAS-6XGFP*. αTi-ltm: control, n=10; *DIP-α^-20F^* n=10; *DIP-α^-50F^*, n=10; *DIP-α^Null^* n=10; αTi-tadm: control, n=12; *DIP-α^-20F^* n=10; *DIP-α^-50F^*n=11; *DIP-α^Null^* n=11. **E**. Quantification of total branch length of control and *DIP-α* mutant alleles over *DIP-α-T2A-Gal4>>20XUAS-6XGFP*. αTi-ltm: control, n=10; *DIP-α^-20F^* n=10; *DIP-α^-50F^* n=10; *DIP-α^Null^* n=10; αTi-tadm: control, n=12; *DIP-α^-20F^* n=10; *DIP-α^-50F^* n=11; *DIP-α^null^* n=10. For all graphs statistical significance was determined using an unpaired nonparametric two tailed Mann-Whitney test. Error bars represent mean with 95% confidence intervals. ns= no statistical difference. *p<0.05 **p<0.01 ***p<0.001 ****p<0.0001.

**Figure 3.**
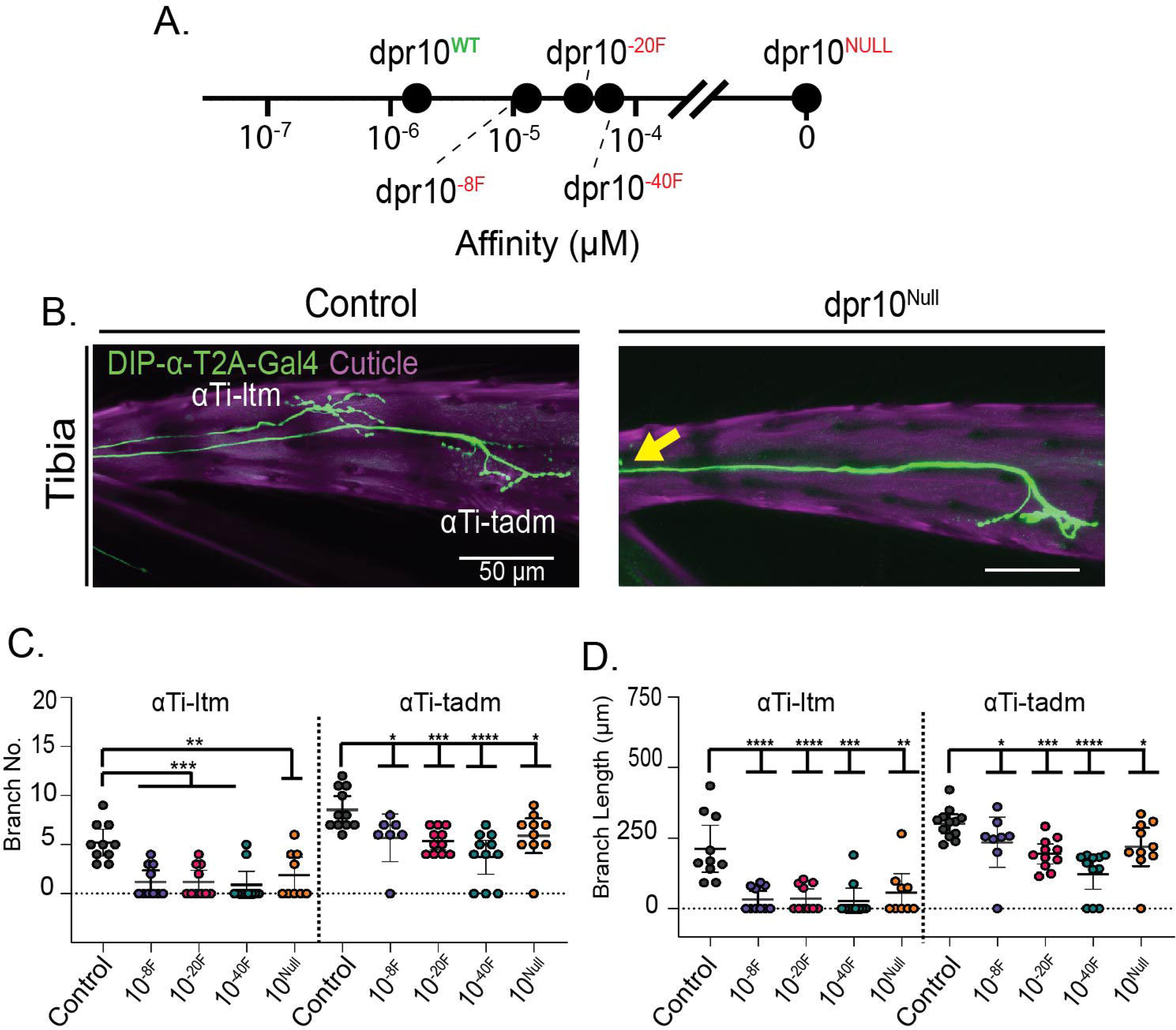
Analysis of *dpr10* alleles with reduced affinity. **A.** Affinity range of *dpr10* mutants in log scale. **B.** Representative images of *yw/DIP-α-T2A-Gal4>>20XUAS-6XGFP* (control) and *yw/DIP-α-T2A-Gal4;dpr6^-^dpr10^-^//>>20XUAS-6XGFP* tibia leg segments. **C**. Quantification of terminal branch number of *dpr10* mutants, *yw/DIP-α-T2A-Gal4; dpr6^-^dpr10^affinity^ ^mutant^//>>20XUAS-6XGFP*. αTi-ltm: control, n=10; *dpr10^-8F^* n=10; *dpr10^-20F^* n=10; *dpr10^-40F^* n=10; *dpr10^Null^* n=9; αTi-tadm: control, n=12; *dpr10^-8F^* n=8; *dpr10^-20F^* n=11; *dpr10^-40F^* n=11; *dpr10^Null^* n=10. **D**. Quantification of total branch length for control and *yw/DIP-α-T2A-Gal4; dpr6^-^ dpr10^affinity^ ^mutant^//>>20XUAS-6XGFP* .αTi-ltm: control, n=10; *dpr10^-8F^* n=10; *dpr10^-20F^* n=10; *dpr10^-40F^* n=10; *dpr10^Null^* n=9; αTi-tadm: control, n=12; *dpr10^-8F^* n=8; *dpr10^-20F^* n=11; *dpr10^-40F^* n=11; *dpr10^Null^* n=10. The control samples in Figure 2D and E were the same because all experiments were done at the same time. For all graphs statistical significance was determined using an unpaired nonparametric two tailed Mann-Whitney test. Error bars represent mean with 95% confidence intervals. ns= no statistical difference. *p<0.05 **p<0.01 ***p<0.001 ****p<0.0001.

Compared to controls, adult animals with *DIP-α* decreased affinity alleles (*DIP-α^-^*^20F^ and *DIP-α^-50F^*) show phenotypes similar to *DIP-α* null animals, where αMN axons fail to arborize onto their muscle target and have no terminal branches (Figure 2B). Unfortunately, using *DIP-α-T2A-Gal4,* we are unable to visualize the αFe-ltm, because even in *DIP-α-T2A-Gal4/+* animals αFe-ltm fails to arborize ∼50% of the time^27^. However, using *A8-Gal4* to directly compare all three αMNs we find that the three αMNs have different expressivities in the absence of *DIP-α* or *dpr10*: αFe-ltm completely fails to arborize, αTi-ltm terminal branches are reduced or lost in about ∼70-80% of animals, and αTi-tadm is rarely affected^27^. In addition to these observations with null alleles, we also find there is a graded affinity requirement for each of the αMNs. For instance, we find that for αFe-ltm there is less arborization for the *DIP-^-^*^50F^ allele compared to the α allele, suggesting that the affinity required to stabilize the branch is within this range (Figure S1). αTi-ltm also fails to arborize although it seems that the expressivity is not as strong as in the αFe-ltm (Figure S1B and C). Finally, although the αTi-tadm rarely shows a complete loss of arborization, both branch number and branch length are reduced (Figure 2D and Figure S2B). Together, these data reveal that the three αMNs all require different minimal affinities to properly arborize onto their target muscles.

### Reducing Dpr10 affinity to DIP-**α** impedes arborization

Considering that we see *DIP-α^-^*^20F^ flies lose the terminal branches of αFe-ltm, we wondered whether we would see a similar loss with the *dpr10^-8F^* allele. As with the *DIP-α* alleles, we examined the *dpr10* alleles using both *DIP-α-T2A-GAL4* and *A8-Gal4* as readouts of αMN morphology. Using the *A8-Gal4* as a readout, we find that for *dpr10^-8F^*, αFe-ltm branches are not completely absent but there is a decrease in branch number and branch length, while the -20F allele of *dpr10* results in the complete loss of branches in this MN (Figure S1D and F). Furthermore, we find that for αTi-ltm the complete absence to arborize occurs at a lower frequency compared to αFe-ltm for all *dpr10* alleles, indicating that the αFe-ltm has a higher affinity requirement than the αTi-ltm (Figure S1). However, using *DIP-α-T2A-Gal4*, αTi-ltm fails to arborize in most *dpr10^-8F^* animals. This discrepancy may be explained because *DIP-α-T2A-Gal4* is a strong hypomorph of *DIP-α* ^27^, possibly resulting in lower levels of DIP-α and a compromised interaction between αMNs and muscles. For αTi-tadm there is a slight decrease in branch number and length in both readouts and in both cases the αTi-tadm arborization phenotype is less strong than with αTi-ltm (Figure 2C and D, Figure S1D and F).

Together, these results suggest that the affinity requirement for the arborization of αFe-ltm is higher than the αTi-ltm, consistent with the differences observed in expressivity for *DIP-α* and *dpr10* null alleles. αTi-tadm is rarely fully lost in either *DIP-α* or *dpr10* null alleles indicating that it relies on additional mechanisms to achieve its proper branch number and length. Finally, these results indicate that varying the affinity from the postsynaptic target has the same effects as varying the affinity from the presynaptic αMN.

### Increasing affinity also impedes **α**MN arborization

Given the loss of arborization when the DIP-α::Dpr10 affinity was reduced, we considered the possibility that increasing the affinity might result in more and/or longer branches. To address this question we used the following *DIP-α* and *dpr10* mutants: *DIP-α*^G74A^ (K_D_ = 0.90 μM, *DIP-α*^+*2F*^) and *dpr10*^Q142M^ (K_D_ = 0.19 μM, *dpr10*^+10F^). In addition, we also analyzed the combination of *DIP-α^+2F^* and *dpr10* ^+10F^, which results in an increased affinity of +15F (*DIP-α & dpr10^+15F^*; K_D_ = 0.10 μM) (Figure 4A). We find that in *DIP-α^+2F^* animals, αMN branch number and length is not affected (Figure 4C). Surprisingly, in *dpr10*^+10F^ flies, αTi-ltm and αTi-tadm branch number and branch length were on average shorter than in control flies (Figure 4C and D). In *DIP-α & dpr10^+15F^* flies, some αTi-ltm and αTi-tadm axons completely fail to arborize. Together, these results show that increasing affinity, like reducing affinity, also adversely affects terminal branch arborization. We note however that using *A8-Gal4* as the readout, the lack of arborization phenotype is not as strong as in the experiments done with the *DIP-α-T2A-Gal4*, even in a sensitized background with only one functional copy of *DIP-α* (*DIP-α & dpr10^+^*^15^ *^Null^*). Nevertheless, there is a reduction in branch number and branch length for both the αFe-ltm and αTi-ltm in this genotype (Figure S2). Together, these experiments indicate that increasing the affinity between DIP-α and Dpr10 only has strong effects when the affinity is above a certain threshold, and that the penetrance of this effect depends on the status of the *DIP-α* locus (see Discussion).

**Figure 4.**
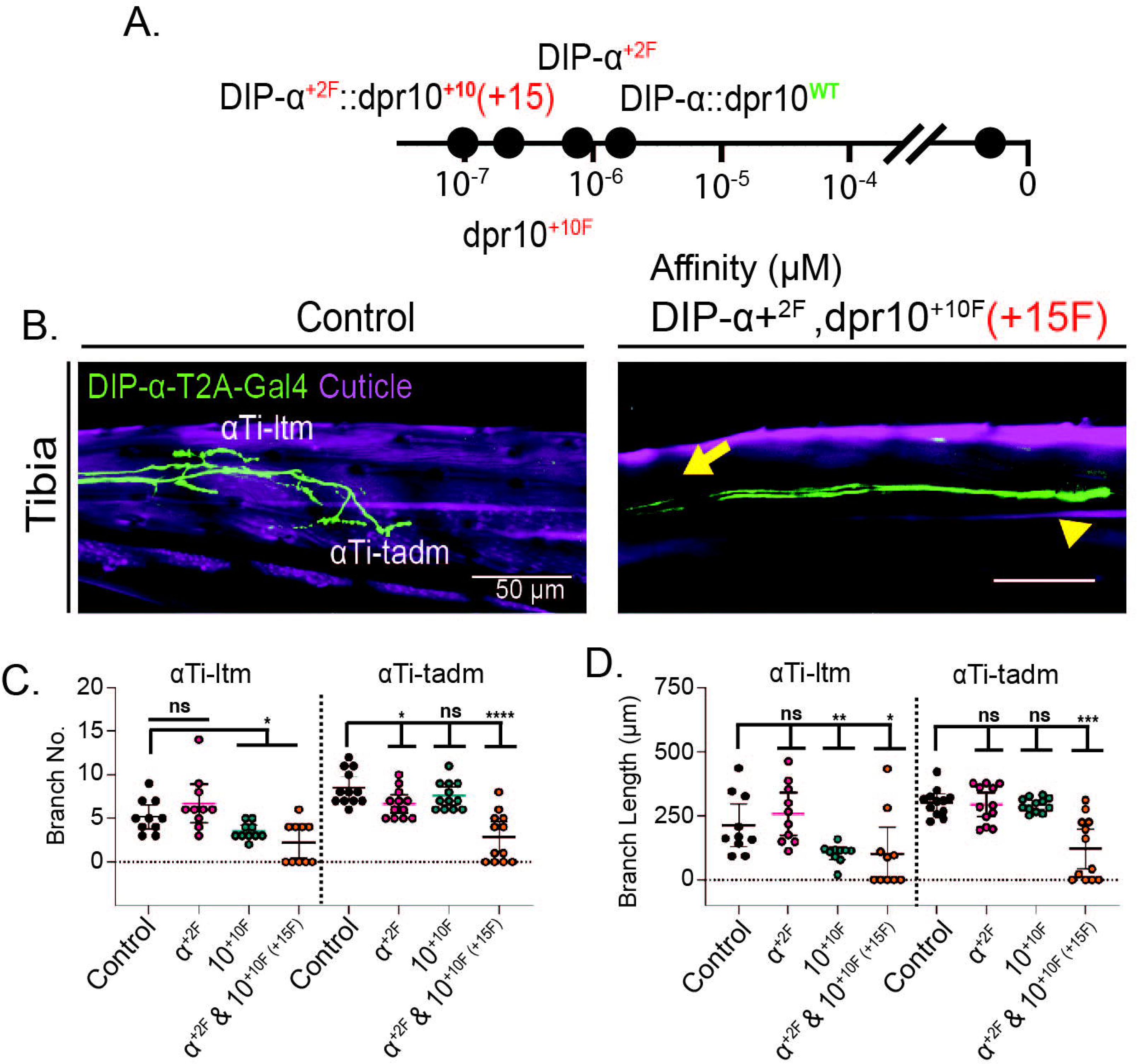
Analysis of increased affinity genotypes. **A.** Affinity range of increased affinity mutants in log scale. **B.** Representative images of *yw/DIP-α-T2A-Gal4>>20XUAS-6XGFP* (control) and *DIP-α^G74A^/DIP-α-T2A-Gal4;dpr6^-^dpr10^Q142M^//>>20XUAS-6XGFP (+15F)* tibia segments. **C**. Quantification of terminal branch number of *dpr10* mutants, *DIP-α^G74A^/DIP-α-T2A-Gal4//>>20XUAS-6XGFP(+2F)*; *yw/DIP-α-T2A-Gal4; dpr6^-^dpr10^Q142M^//>>20XUAS-6XGFP(+10F)*; and *DIP-α^G74A^/DIP-α-T2A-Gal4; dpr6^-^dpr10^Q142M^//>>20XUAS-6XGFP(+15F*). αTi-ltm: control, n=10; *DIP-α^+2F^* n=10; *dpr10^+10F^* n=10; *DIP-α^+2F^* & *dpr10^+10F^* n=10; αTi-tadm: control, n=12; *DIP-α^+2F^* n=12; *dpr10^+10F^* n=12; *DIP-α^+2F^* & *dpr10^+10F^* n=12. **D**. Quantification of total branch length in control and *DIP-α^G74A^/DIP-α-T2A-Gal4//>>20XUAS-6XGFP(+2F); yw/DIP-α-T2A-Gal4; dpr6^-^dpr10^Q142M^//>>20XUAS-6XGFP(+10F);* and *DIP-α^G74A^/DIP-α-T2A-Gal4; dpr6^-^dpr10^Q142M^//>>20XUAS-6XGFP (+15F)* flies. αTi-ltm: control, n=10; *DIP-α^+2F^* n=10; *dpr10^+10F^* n=10; *DIP-α^+2F^* & *dpr10^+10F^* n=10; αTi-tadm: control, n=12; *DIP-α^+2F^* n=12; *dpr10^+10F^* n=12 *DIP-α^+2F^* & *dpr10^+10F^* n=12. The control samples for Figure 2D and E were the same because all experiments were done at the same time. For all graphs statistical significance was determined using an unpaired nonparametric two tailed Mann-Whitney test. Error bars represent mean with 95% confidence intervals. ns= no statistical difference. *p<0.05 **p<0.01 ***p<0.001 ****p<0.0001.

To determine if αMNs with reduced arborization are able to make synapses, we expressed the marker Bruchpilot (Brp-short::mStraw^38^) in *DIP-^+15F^* and control flies. Although we were not able get single bouton resolution of Brp puncta, we are nevertheless able to see that in *DIP-α & dpr10^+15F^*flies, αTi-ltm and αTi-tadm have Brp puncta, suggesting that they assemble synapses (Figure S3).

### Increasing affinity prevents the stabilization of developing **α**MNs

It is possible that *DIP-α & dpr10^+15F^*αMNs do not stabilize on their muscle targets either because 1) they never send out tertiary branches or 2) they send out branches but they are not capable of stabilizing on their targets. To test this, we took time lapse videos and compared control to *DIP-α & dpr10^+15F^* flies, imaged from ∼25-80 hrs APF, the time window when αMNs form their secondary and terminal branches and stabilize on their muscle targets^27^ (Video 2). In control animals, at ∼25 hrs APF both αTi-ltm and αTi-tadm have already reached their muscle targets. Between 35-55 hrs APF the axons will branch and extend filopodia in the general area of their muscle targets and then at ∼60 hrs APF some filopodia will stabilize. In *DIP-α & dpr10^+15F^*animals, both αTi-ltm and αTi-tadm reach their muscle targets successfully at 24 hrs APF. Very few differences can be seen in the next 24 hours. However, at 60 hrs APF the tertiary branches that have extended fail to stabilize and instead retract resulting in the phenotype that we see in the adult. Interestingly, this is similar to what has been described for *DIP-α* nulls (Figure 5A). These results indicate that αMNs can initially “recognize” the correct muscle target and send out filopodia to the correct myoblast cluster but fail to maintain the interaction long enough for the axons to fully arborize.

**Figure 5.**
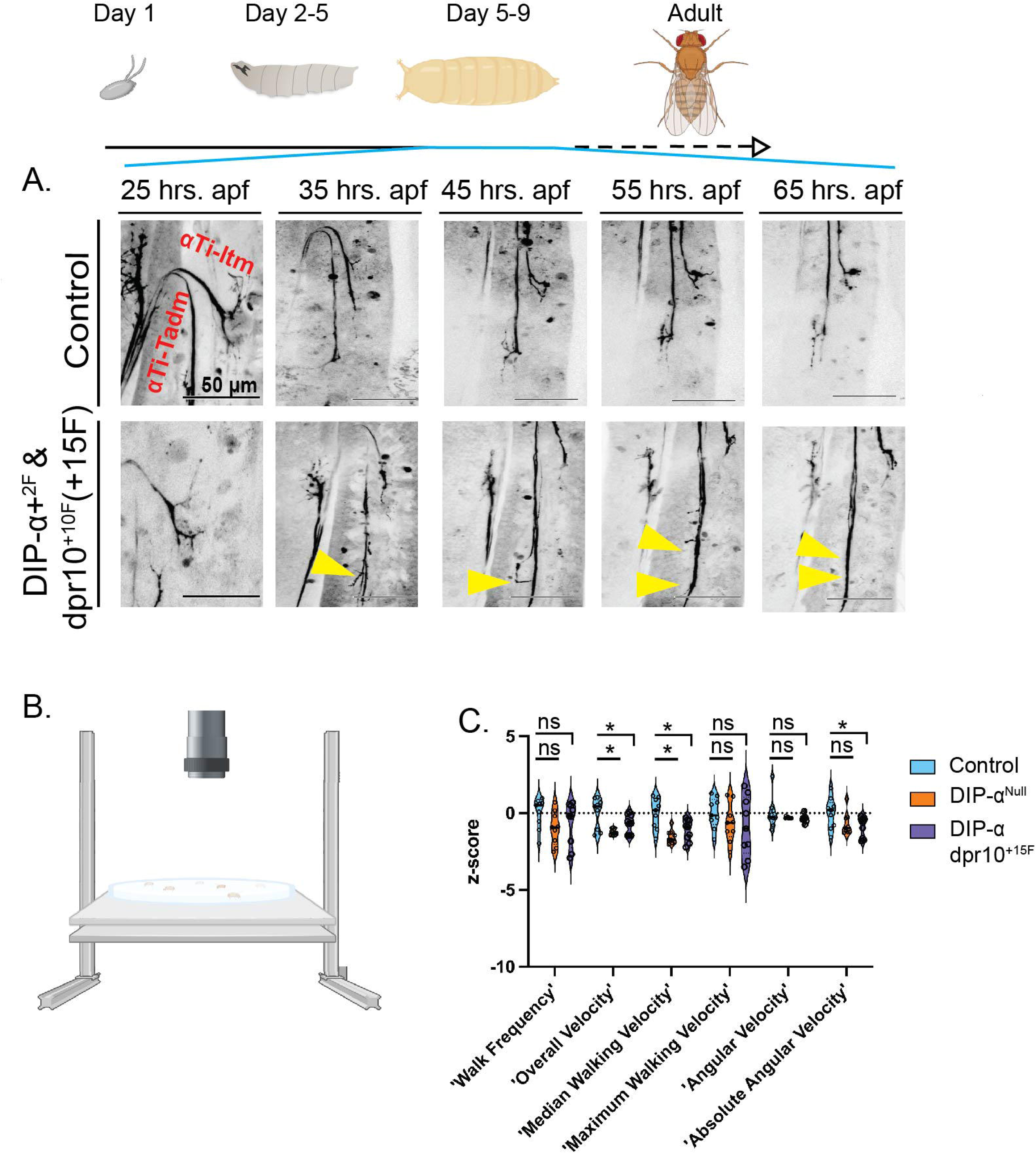
Live imaging and locomotion analysis. **A.** Still images of a timelapse video (see Video 2) of *yw/DIP-α-T2A-Gal4>>20XUAS-6XGFP* pupa (top row) and *DIP-α^G74A^/DIP-α-T2A-Gal4;dpr6^-^dpr10^Q142M^//>>20XUAS-6XGFP (+15F)* (bottom row) flies. Yellow arrowheads point to small filopodia that aberrantly extend but ultimately retract in *DIP-α^G74A^/DIP-α-T2A-Gal4; dpr6^-^dpr10^Q142M^* MNs. **B.** Free walking arena schematic. **C**. Walking parameters measured in control (*yw*, *20XUAS-6XGFP*); *DIP-α^Null^*, *20XUAS-6XGFP*, and +15F gain-of-affinity *DIP-α^G74A^//20XUAS-6XGFP;dpr6^-^dpr10^Q142M^(+15)//* genotypes. Control n=10; *DIP-α^Null^* n=10; *DIP-α & dpr10^+15F^* n=10. *p<0.05

### Changing affinity leads to behavioral defects

The three αMNs directly target the Fe-ltm, Ti-ltm, and Ti-tadm muscles^27^. It is plausible that locomotion defects might occur when the terminal branches in these MNs fail to properly arborize. Previously our lab established a free walking behavioral assay that reports a dozen different walking parameters^39^ (Figure 5B). We compared both *DIP-α^Null^* and *DIP-α & dpr10^+15F^* to control animals because these alleles are highly penetrant. Both *DIP-α^Null^* and *DIP-α & dpr10^+15F^*flies have a decreased overall velocity and median walking velocity. Furthermore, *DIP-α & dpr10^+15F^*flies also have a reduced absolute angular velocity compared to controls (Figure 5C). Although, it is possible that these effects are the result of other, non-MN *DIP-α*-expressing neurons elsewhere in the nervous system, they demonstrate that changing the affinity between DIP-α::Dpr10 adversely affects locomotion.

## Discussion

Here we use single neuron morphology measurements to understand the importance of affinity between two CAMs that are required to maintain contacts between MNs and muscles in the Drosophila leg neuromuscular system^37^. We address this question using a series of knock in alleles of both *DIP-α* and *dpr10* that change the affinity of these proteins, as measured *in vitro*^33^. We show that decreasing the affinity past a certain threshold leads to phenotypes similar to the null phenotype^27^. Surprisingly, we also find that increasing the affinity past a certain threshold also reduces the amount of axonal arborization. We also show via live imaging that axons in flies with high affinity alleles are able to extend filopodia to the correct muscle target but ultimately fail to stabilize and arborize. Together, these data show that a specific affinity window for these interacting CAMs is needed for proper arborization.

The loss of branching in the high affinity mutants was initially surprising. Our lab has recently shown that *dpr10* is also expressed by αMNs and that removing MN-expressed *dpr10* leads to abnormally long branches in both the αTi-ltm and αTi-tadm^40^. Therefore, a possible mechanism that contributes to the phenotype seen in the *DIP-α & dpr10^+15F^* mutant is that the higher affinity interaction between DIP-α and Dpr10 isoccurring in *cis* (in MNs), which reduces DIP-α’s ability to interact with Dpr10 in *trans* in muscle cells. This could explain why the filopodia still target the correct muscles but are not able to “anchor” long enough to generate stable terminal branches. Dpr10 is also expressed by other MNs in addition to the αMNs. Therefore, another explanation is that DIP-α in αMNs is interacting with Dpr10 present in other MNs. However, this seems less likely as we would expect the filopodia to become less mobile in the *DIP-α & dpr10^+15F^* mutant due to more and stronger interneuronal contacts.

Another interesting observation is that, like the time lapse videos of *DIP-α null*^27^, the live imaging videos of *DIP-α & dpr10^+15F^* mutants reveal that filopodia still extend out to their correct muscle target but unable to make stable contacts with their target and then retract (Figure 5A and Video 2). This reinforces the idea that DIP-α::Dpr10 *trans* interactions are needed to stabilize terminal branches, and not for axon guidance or synaptic specificity. Further, this suggests that there must be additional mechanisms that help MN axons recognize their targets. We propose that the proper DIP-α::Dpr10 affinity range is needed to stabilize the contact long enough to recruit more DIP-α::dpr10 *trans* interactions and keep the axon arborizing.

It is not clear why the phenotype is stronger in *DIP-α & dpr10^+15F^/DIP-α-T2A-Gal4* animals compared to *DIP-α & dpr10^+15F^* /*DIP-* ^Null^ animals using *A8-Gal4*, when theoretically the amount of DIP-α should be the same in both genotypes. It is possible that a truncated DIP-α protein generated by the *DIP-α-T2A-Gal4* allele is further sensitizing the phenotype. One way to test this is to overexpress Dpr10 in αMNs and/or in muscles in *DIP-α & dpr10^+15F^*/*DIP-* ^Null^ flies and see whether we obtain similar phenotypes to those seen in *DIP-α & dpr10^+15F^/DIP-α-T2A-Gal4* flies. This would also allow us to determine whether the phenotype is due to differences in levels in the MNs or the muscles. We have shown that overexpressing *dpr10* in muscle does not have a phenotype but that overexpressing *dpr10* and *dpr6* in αMNs leads to *cis*-inhibition and a reduction in aborization^40^. Therefore, it would be interesting to see if *DIP-α & dpr10^+15F^*/*DIP-α* ^Null^; *A8-Gal4>>UAS-dpr10* flies have the same phenotype and is indeed a result of *cis* inhibition.

Furthermore, it is interesting that different neural systems utilize the same molecules for different purposes. As noted in the Introduction, reducing DIP-α::Dpr10 affinity in the visual system leads to cell survival and synaptic targeting defects. Here, DIP-α::Dpr10 is not required for cell survival nor synapse formation. We propose that DIP-α and Dpr10 interact with different molecules in different contexts to achieve a variety of functions. These differences may also be a consequence of the more hard-wired nature of how MNs are born compared to the more plastic way in which an excess of neurons are born in the visual system, leading some to apoptose.

## Methods

### Dissection and microscopy

For adult leg dissection, flies were first immersed in 70% Ethanol for ∼1 min and rinsed in 0.3% Triton-X in PBS 3X. Abdomen and heads were then removed and legs attached to thorax were fixed overnight at 4°C in 4% paraformaldehyde in 0.3% Triton-X in PBS. The next day, legs and thorax were washed five times in 0.3% Triton-X in PBS then stored in 80% VECTASHIELD (in PBS) overnight at 4°C. Legs were then removed from the thorax and placed onto a glass slide in a drop of VECTASHIELD mounting medium (Vector Labs). 18X18mm coverslips were then placed on top with dental wax on the corners to prevent the coverslip from crushing the legs. 0.5 µm-thick sections in the Z axis were imaged using a Zeiss LSM 800 Confocal Laser Scanning Microscope.

### Motor neuron axon branch quantification and statistical analysis

To quantify leg motor neurons only one T1 leg from each animal was imaged and their neurons traced using the Simple Neuron Tracer from ImageJ^37^. For each neuron, with the exception of the αTi-tadm, the tracing began at the first bifurcation point. For the αTi-tadm, only the most distal branch was traced due to presence of collateral branches which would have potential misrepresented the terminal branch number and branch length. Both total terminal branch number and the sum of cable length values were separately plotted and analyzed using the GraphPad Prism 9.0 software. For all graphs statistical significance was determined using an unpaired nonparametric two tailed Mann-Whitney test. Error bars represent mean with 95% confidence intervals. ns= no statistical difference. *p<0.05 **p<0.01 ***p<0.001 ****p<0.0001.

### Live Imaging of MN development in pupa

Pupae were first staged by picking late (wandering) L3 larva and placing them in a separate vial with a moist tissue, kept at 25°C and then picked 24 hours later. A small window on the left ventral side of the pupal case was made using forceps to expose just the T1 leg. Individual pupae were placed on a glass slide, surrounded by two layers of filter paper dampened with distilled water. A 5 ul drop of distilled water was placed at the center of a glass coverslip (N-1.5) and placed exactly over the exposed T1 leg. Petroleum jelly surrounding the filter paper was used to seal the space between the coverslip and the glass slide to retain humidity. Samples were imaged on a Zeiss LSM 800 microscope, 25X objective, with a 10 min interval between each z-stack series. Videos were generated using the FIJI software^41^ at five frames per second.

### Fly arena experiments

The fly arena setup was constructed by a previous member of the lab and described in Howard et al, 2009^39^. Briefly, the polycarbonate plastic arena sits in an aluminum plate, which maintains a level surface and is covered with a circular acrylic disc that has a small hole for mouth pipetting in flies. The inside of the lid is freshly coated with a thin layer of fluon (Amazon, B00UJH12A) to prevent flies from walking on the ceiling. Videos were recorded with a Point Grey Blackfly Mono USB3 camera fitted with a Tamron ½” F/1.2 ER C-mount lens (B&H photo) was mounted above the arena and connected to a computer running Ubuntu 20.04 LTS. A Kodax 3x3” 89B Opaque IR filter (B&H photo) was placed in front of the camera detector to allow for detection of IR but not visible light. Backlighting was provided by a plate of LEDs arranged in parallel rows underneath the arena. An acrylic diffuser was placed in front of the camera detector to allow for detection of IR but not visible light. Infrared LEDs were acquired from ledlightsworld.com (SMD3528-300). The IR lights were set at 100% brightness.

### Data Acquisition

For recordings, 3-5 day old mated females were kept in a humidity and temperature controlled incubator with a 12 hr day/night. All behavior videos were recorded between the morning activity peak time (9am-12pm). Prior to the recording, the arena was leveled, the lid cleaned and layered with fluon to prevent the flies from walking on the lid. Each recorded video had 10 flies. Videos were taken on multiple days and pooled for the final dataset (N = 30). For every recording session, flies were mouth pipetted into the arena through the small hole in the lid and the lid was slid to move the hole out of the camera view. A blackout curtain was used to cover the arena to protect from any contaminating light. All videos were recorded at a rate of 30 frames per second and stored in a compressed “fly format” using software written by Andrew Straw at the University of Frieburg. Videos were tracked using the Matlab based FlyTracker software from the Caltech vision lab^42^. Before tracking, pixel to mm conversion was calibrated using the inbuilt GUI of the tracker software and a single calibration file was generated for all the videos recorded on the same day. Background model and thresholds were adjusted to provide optimal recognition of animals and were not standardized between recording sessions. All analysis on data from arena experiments was performed in MATLAB using scripts written by Dr. Clare E. Howard and modified by Dr. Sumaira Zamurrad.

### Behavioral classifiers

<u>Walk:</u> Walking frames were defined using a dual threshold Schmitt trigger filter (original script written by Irene Kim – Dickinson lab). Speed thresholds were set at 1 and 2.5 mm/s, and time thresholds were .1 s. Walking frames were also specified to be those in which the fly was not already engaged in a jump.

Stop: Stop frames were classified as any frames where animals were not performing walking or jumping behaviors.

### Parameters

Walk Frequency: the percent of frames classified as walking during the recording period.

Overall Velocity: the median of all velocities over the recording period.

Median Walking Velocity: the median of velocities during all frames when the animal is classified as walking.

Maximum Walking Velocity: the maximum velocity an animal reaches during walking.

<u>Angular velocity:</u> the median value of angular velocity. This parameter takes into account directionality of turning.

<u>Absolute Angular Velocity:</u> the median of the absolute value of angular velocities. This parameter does not take into account directionality of turning.

Walking bout duration: the length of each bout was calculated, and the median of all bout lengths was taken for each animal.

Stop bout number and duration: calculated as for walking bouts.

For each animal, the value of each parameter was defined as described above for a ten-minute walking period in the dark. Comparisons with control were drawn with Kruskall-Wallis (for non-normally distributed data) analysis.

## Acknowledgements

This work was funded by a grant from NINDS, NS070644, to R.S.M. We thank Drs. Honig, Shapiro, Zinn, and Zipursky for sharing data and ideas in the early stages of this work and Dr. L. Venkatasubramanian for training and advice in the early phase of this project.

## Author contributions

D.L. executed all of these experiments, including the statistical analyses, with help from K.R. S.Z. helped with the behavioral analysis. S.X. provided the knock-in alleles and additional information regarding these chromosomes. D.L. wrote the initial draft. R.S.M. conceptualized the study, provided feedback, and edited the manuscript.

## Supplementary Figure Legends

**Figure S1. Analysis of decreased affinity mutants using *A8-Gal4* as readout.**

**A.** Representative femur and tibia images of *yw, A8-Gal4>>20XUAS-6XGFP* (control) and *DIP-α ^Null^, A8-Gal4>>20XUAS-6XGFP* femur and tibia leg segments.
**B.** Quantification of terminal branch number of *DIP-α* mutants; αFe-ltm: control n=8; *DIP-α^-20F^, A8-Gal4* n=10; *DIP-α^-50F^, A8-Gal4* n=10; *DIP-α^Null^, A8-Gal4* n=10; αTi-ltm: control n=9; *DIP-α^-20F^, A8-Gal4* n=9; *DIP-α^-50F^, A8-Gal4* n=10; *DIP-α^Null^, A8-Gal4* n=8; αTi-tadm: control n=9; *DIP-α ^-20F^, A8-Gal4* n=10; *DIP-α^-50F^, A8-Gal4* n=9; *DIP-α ^Null^, A8-Gal4* n=10.
**C.** Quantification of total branch length of control and *DIP-α* mutants. αFe-ltm: control α , *A8-Gal4* n=10; *DIP-α* , *A8-Gal4* n=10; *DIP-α* , *A8-Gal4* n=10; αTi-ltm: control n=9; *DIP-α^-20F^, A8-Gal4* n=9; *DIP-α^-50F^, A8-Gal4* n=10; *DIP-α^Null^, A8-Gal4* n=9; αTi-tadm: control n=9; *DIP-α^-20F^, A8-Gal4* n=10; *DIP-α^-50F^, A8-Gal4* n=9; *DIP-^Null^,A8-Gal4* n=10.
**D.** Representative images of *yw, A8-Gal4>>20XUAS-6XGFP* (control) and *dpr6^-^,dpr10^-^,A8-Gal4>>20XUAS-6XGFP* femur and tibia leg segments.
**E.** Quantification of terminal branch number of *dpr10* mutants. αFe-ltm: control n= 11; *dpr6^-^,dpr10^-8F^, A8-Gal4* n= 9*; dpr6^-^,dpr10^-20F^, A8-Gal4* n=10; *dpr6^-^,dpr10^-40F^, A8-Gal4* n=11; *dpr6^-^, dpr10^Null^, A8-Gal4* n= 12; αTi-ltm: Control n= 11; *dpr6^-^,dpr10^-8F^, A8-Gal4* n= 9; *dpr6^-^,dpr10^-20F^, A8-Gal4* n=10; *dpr6^-^,dpr10^-40F^, A8-Gal4* n=11; *dpr6^-^, dpr10^Null^, A8-Gal4* n= 12; αTi-tadm: Control n= 11; *dpr6^-^,dpr10^-8F^, A8-Gal4* n= 9; *dpr6^-^,dpr10^-20F^, A8-Gal4* n=10; *dpr6^-^,dpr10^-40F^, A8-Gal4* n=10; *dpr6^-^, dpr10^Null^, A8-Gal4* n=12.
**F.** Quantification of total branch length of *dpr10* mutants. αFe-ltm: control n=11; *dpr6^-^ ,dpr10^-8F^, A8-Gal4* n=9; *dpr6^-^,dpr10^-20F^, A8-Gal4* n=10; *dpr6^-^,dpr10^-40F^, A8-Gal4* n=11; *dpr6^-^, dpr10^Null^, A8-Gal4* n=12; αTi-ltm: control n=11; *dpr6^-^,dpr10^-8F^, A8-Gal4* n=9; *dpr6^-^,dpr10^-20F^, A8-Gal4* n=10; *dpr6^-^,dpr10^-40F^, A8-Gal4* n=11; *dpr6^-^, dpr10^Null^, A8-Gal4* n=12; αTi-tadm: control n=11; *dpr6^-^,dpr10^-8F^, A8-Gal4* n=9; *dpr6^-^,dpr10^-20F^, A8-Gal4* n=10; *dpr6^-^,dpr10^-40F^, A8-Gal4* n=11; *dpr6^-^, dpr10^Null^, A8-Gal4* n=12. For all graphs statistical significance was determined using an unpaired nonparametric two tailed Mann-Whitney test. Error bars represent mean with 95% confidence intervals. ns= no statistical difference. *p<0.05 **p<0.01 ***p<0.001 ****p<0.0001. All quantifications done with *20XUAS-6XGFP*.

**Figure S2. Analysis of increased affinity mutants with A8-Gal4 as the readout.**

**A.** Representative images of control, *yw, DIP-α^Null^, A8-Gal4, dpr6 dpr10^Q^*^142^*^M^ heterozygous>>20XUAS-6XGFP* and *DIP-α^G74A^, DIP-α^Null^, A8-Gal4, dpr6 dpr10^Q142M^* >>*20XUAS-6XGFP* homozygous femur and tibia leg segments.
**B.** Quantification of terminal branch number of control, *yw, DIP-α^Null^, A8-Gal4, dpr6-dpr10^Q142M^* heterozygous*>>20X-6XGFP* and *DIP-α*^G74A^, *DIP-α^Null^, A8-Gal4, dpr6-dpr10^Q142M^* homozygous genotypes. αFe-ltm: control n=11; *DIP-α^+2^, DIP-α^Null^*, *dpr6-dpr10^+10^*n=13; αTi-ltm: control n=10; *DIP-α^+2^, DIP-α^Null^, dpr6-dpr10^+10^* n=13; αTi-tadm: control n=11; *DIP-α^+2^, DIP-α^Null^, dpr6-dpr10^+10^* n=13.
**C.** Quantification of total branch length of control *yw, DIP-α^Null^, A8-Gal4, dpr6-dpr10^Q142M^*heterozygous*>>20X-6XGFP* and *DIP-α^G74A^/ DIP-α^Null^, A8-Gal4, dpr6-dpr10^Q142M^* homozygous genotypes. αFe-ltm: control n=11, *DIP-α^+2^/DIP-α^Null^, dpr6-dpr10^+10^* n=13; αTi-ltm: control n=10; *DIP-α^+2^/ DIP-α^Null^, dpr6-dpr10^+10^* n=14; αTi-tadm: control n=11; *DIP-α^+2^/ DIP-α^Null^, dpr6-dpr10^+10^* n=13. For all graphs statistical significance was determined using an unpaired nonparametric two tailed Mann-Whitney test. Error bars represent mean with 95% confidence intervals. ns= no statistical difference. *p<0.05 **p<0.01 ***p<0.001 ****p<0.0001. All quantifications were done with *20XUAS-6XGFP*.

**Figure S3. Truncated axon termini have boutons**

**A.** Representative images of tibia leg segments from control, *yw*, *DIP-α-T2A-Gal4, dpr6-dpr10^Q142M^ heterozygous>>20XUAS-6XGFP, UAS-brp-short::mStraw* and *DIP-α^G74A^, DIP-α-T2A-Gal4, dpr6-dpr10^Q142M^ homozygous>>20XUAS-6XGFP, UAS-brp-short::mStraw* flies.
**B.** High magnification images of boxed regions in **A** for the control. **B:** αTi-ltm. **B’:** αTi-tadm.
**C.** High magnification images of boxed regions in **A** for the +15F genotype. **C:** αTi-ltm.

**C’:** αTi-tadm.

**Figure S4. Leg myoblast fusion during pupal development.**

Still images of Mef2 expressing leg myoblasts during pupal development. Arrowhead points to Ti-ltm connecting to femur muscles. Arrows point to two femur muscle groups segregating during development.

## Video legends

**Video 1**: https://www.dropbox.com/scl/fi/idapupdlq2gkcb033upzg/Video_1.mov?rlkey=fjwstxhhs4h4wepbkzbxhrlqt&dl=0

*DIP-α-T2A-Gal4>>20XUAS-6XGFP,Mef2-QUAS>>10XQUAS-6xmCherry*.

Green=αMNs; Magenta=myoblasts

Video 2:

https://www.dropbox.com/scl/fi/9wfaszkqwem6eyw33rnzc/Video_2.mov?rlkey=ln4xzlqn9t7ttf5ile6vtdg3g&dl=0

*Left: yw/DIP-α-T2A-Gal4>>20XUAS-6XGFP; Right: DIP-α^+2^/DIP-α-T2A- Gal4;dpr6,dpr10^+10^ (+15)>>20XUAS-X6GFP*.

**Video S1**: https://www.dropbox.com/scl/fi/t1hjvkovek7n1xsteoj4z/Video_S1.mov?rlkey=zotp832ddptw9i3aarqq8x058&dl=0

*VGlut-LexA>>13XLexAop-myr::GFP, Mef2-Gal4>10XUAS-myr::tdTomato*

