## Supplementary figures and images for "A critical affinity window for IgSF proteins DIP-α and Dpr10 is required for proper motor neuron arborization"

### Figure S1.jpg

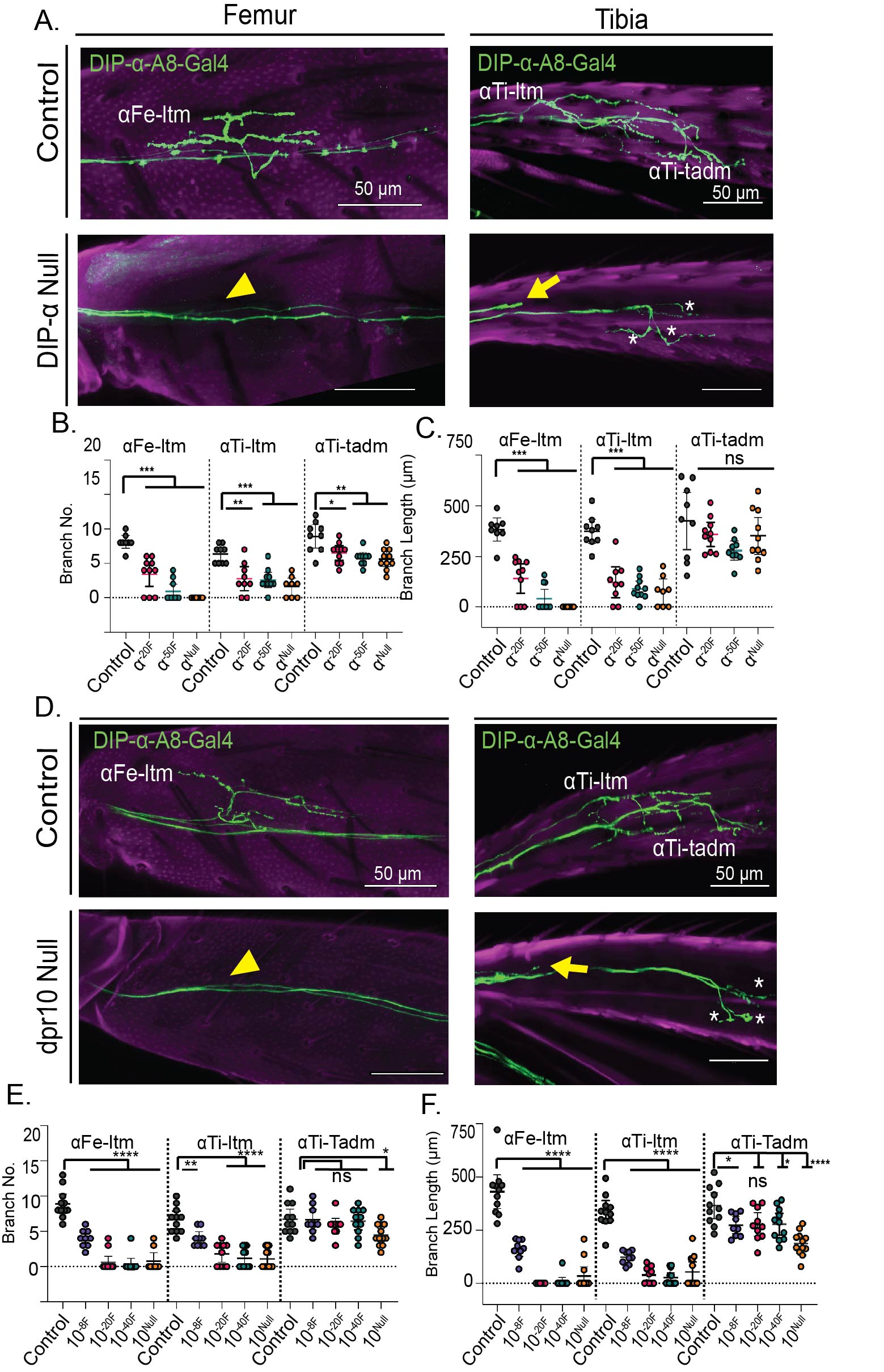

### Figure S2.jpg

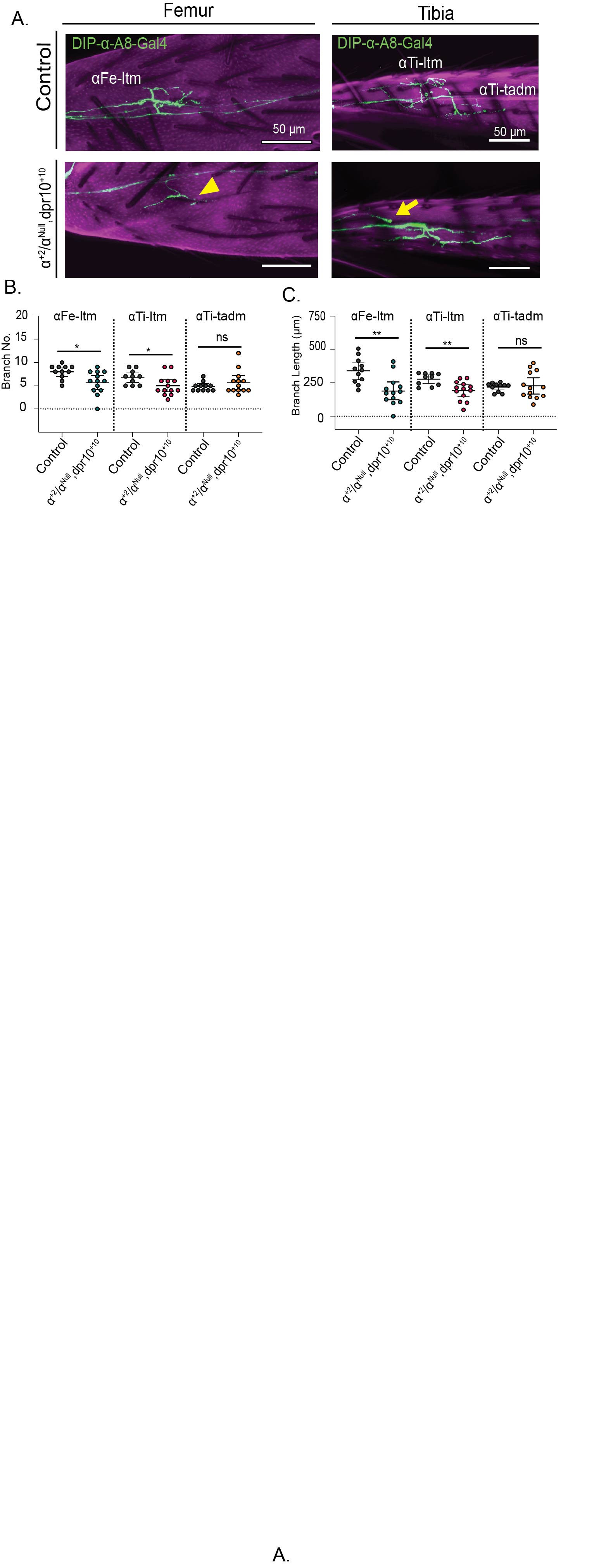

### Figure S3.jpg

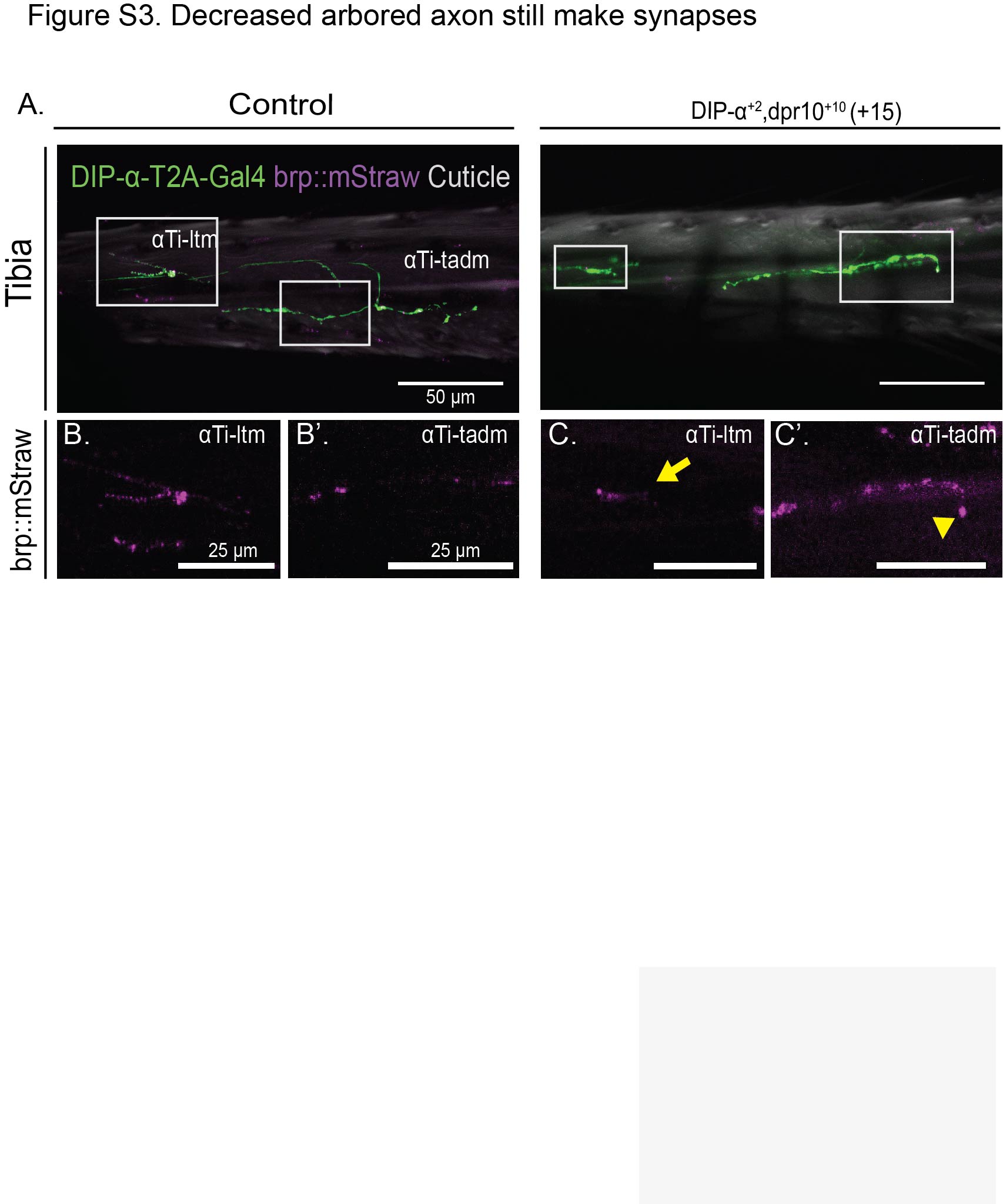
